# Transient inhibition to light explains stronger V1 responses to dark stimuli

**DOI:** 10.1101/2021.11.21.469446

**Authors:** David St-Amand, Curtis L. Baker

**Affiliations:** McGill Vision Research Unit, Department of Ophthalmology & Visual Sciences, McGill University, Montreal, Quebec, Canada, H3G 1A4.

## Abstract

Neurons in the primary visual cortex (V1) receive excitation and inhibition from distinct parallel pathways processing lightness (ON) and darkness (OFF). V1 neurons overall respond more strongly to dark than light stimuli, consistent with a preponderance of darker regions in natural images, as well as human psychophysics. However, it has been unclear whether this “dark-dominance” is due to more excitation from the OFF pathway or more inhibition from the ON pathway. To understand the mechanisms behind dark-dominance, we record electrophysiological responses of individual simple-type V1 neurons to natural image stimuli and then train biologically inspired convolutional neural networks to predict the neurons’ responses. Analyzing a sample of 74 neurons (in anesthetized, paralyzed cats) has revealed their responses to be more driven by dark than light stimuli, consistent with previous investigations. We show that this asymmetry is predominantly due to slower inhibition to dark stimuli rather than to stronger excitation from the thalamocortical OFF pathway. Consistent with dark-dominant neurons having faster responses than light-dominant neurons, we find dark-dominance to solely occur in the early latencies of neurons’ responses. Neurons that are strongly dark-dominated also tend to be less orientation selective. This novel approach gives us new insight into the dark-dominance phenomenon and provides an avenue to address new questions about excitatory and inhibitory integration in cortical neurons.

**Significance:** Neurons in the early visual cortex respond on average more strongly to dark than to light stimuli, but the mechanisms behind this bias have been unclear. Here we address this issue by combining single-unit electrophysiology with a novel machine learning model to analyze neurons’ responses to natural image stimuli in primary visual cortex. Using these techniques, we find slower inhibition to light than to dark stimuli to be the leading mechanism behind stronger dark responses. This slower inhibition to light might help explain other empirical findings, such as why orientation selectivity is weaker at earlier response latencies. These results demonstrate how imbalances in excitation vs. inhibition can give rise to response asymmetries in cortical neuron responses.

## Introduction

The early thalamocortical visual system is separated into two distinct pathways: an ON pathway which responds more to lighter parts of images and an OFF pathway which encodes darker image regions. Neurons in primary visual cortex (V1) combine inputs from these two pathways, but the nature of this integration is still poorly understood.

V1 neurons evidently receive asymmetrical inputs from the two pathways, since they are on average more responsive to dark than light stimuli (Jin et al., 2008; Yeh et al., 2009), especially at low spatial frequencies (Kremkow et al., 2014; Jansen et al., 2019) and shorter time latencies (Komban et al., 2014). This asymmetry is presumably adaptive due to the preponderance of dark regions in natural images (Ratliff et al., 2010), which is also more pronounced at lower spatial frequencies (Cooper & Norcia, 2015). These asymmetries may influence human perception, since dark stimuli are processed faster and more reliably than light stimuli (Buchner & Baumgartner, 2007; Komban, Alonso & Zaidi, 2011).

There are more OFF than ON excitatory inputs from the lateral geniculate nucleus (LGN) to layer 4 of V1, which could help explain why responses to dark stimuli are stronger in V1 (Jin et al., 2008). However, this does not explain why more dark-dominant neurons are found in layers 2/3 than in layer 4 (Yeh et al., 2009). This discrepancy could be explained by stronger ON than OFF intracortical inhibition within V1 (Taylor, Sedigh-Sarvestani, Vigeland, Palmer, & Contreras, 2018). Hence, whether dark-dominance is mostly due to excitation to dark stimuli or inhibition to light stimuli remains unclear. Here we develop a novel machine learning approach to disambiguate excitation from inhibition in extracellular recordings, which allows us to make quantitative inferences about how cortical neurons integrate ON and OFF inputs.

To better understand how visual stimuli drive V1 responses, we predict the responses of recorded neurons to natural images with a simple, biologically-inspired convolutional neural network. This neural network processes the natural images’ light (ON) and dark (OFF) information in two distinct pathways. The first layer of each pathway consists of a convolution with a parametrized 2D gaussian spatial filter, which represents the responses of LGN neurons (omitting the weaker surrounds; Croner & Kaplan, 1995). The second layer is a linear weighted sum of the excitatory or inhibitory contributions of each pathway, which then sum to provide the model’s output. From these estimated weights, we infer how much excitation and inhibition arises from each pathway, at every spatial location and temporal lag of a V1 cell’s receptive field.

Using this approach, we find the dark-dominance phenomenon in V1 neurons to only occur at the early response latencies. We show these stronger dark responses to be predominantly driven by a lack of inhibition to dark stimuli at early latencies. We also find that this slower inhibition to dark stimuli is associated with less orientation selectivity in neurons’ early responses (Ringach, Hawkin, & Shapley, 1997; Shapley, Hawken & Ringach, 2003). These findings suggest that slower inhibition to dark than to light stimuli plays a crucial role in the dark-dominance found in primary visual cortex.

## Methods

### Animal preparation

Anesthesia in adult cats was induced by isoflurane-oxygen (3–5%) inhalation, followed by intravenous cannulation and bolus injection of propofol (5 mg/kg). Surgical anesthesia was maintained with supplemental doses of propofol. Glycopyrrolate (30 μg) and dexamethasone (1.8 mg) were administered and a tracheal cannula or intubation tube was inserted. Throughout the surgery, body temperature was thermostatically maintained and heart rate was monitored (Vet/Ox Plus 4700).

The animal was then positioned in a stereotaxic apparatus and connected to a ventilator (Ugo Basile 6025). Cortical Area 17 was exposed by a craniotomy (P3/L1) and a small durotomy, and the cortical surface protected with 2% agarose capped with petroleum jelly. Local injections of bupivacaine (0.50%) were administered at all surgical sites. During recording, anesthesia was maintained by infusion of propofol (5.3 mg·kg^-1^·h^-1^), and in addition, remifentanil (initial bolus injection, 1.25 μg·kg^-1^, then infusion, 3.7 µg·kg^-1^·h^-1^) and O_2_/N_2_O (30:70 ratio) delivered through the ventilator. Paralysis was produced with a bolus iv injection of gallamine triethiodide (to effect), followed by infusion (10 mg·kg^−1^·h^−1^). Throughout subsequent recording, expired CO_2_, EEG, ECG, body temperature, blood oxygen, heart rate, and airway pressure were monitored and maintained at appropriate levels. Intramuscular glycopyrrolate (16 μg) and dexamethasone (1.8 mg) were also administered daily.

Corneas were initially protected with topical carboxymethylcellulose (1%) and subsequently with neutral contact lenses. Spectacle lenses were selected with slit retinoscopy to produce emmetropia at 57 cm, and artificial pupils (2.5 mm) were provided. Topical phenylephrine hydrochloride (2.5%) and atropine sulfate (1%), or cyclopentolate (1.0 %) in later experiments, were administered daily.

All animal procedures were approved by the McGill University Animal Care Committee and are in accordance with the guidelines of the Canadian Council on Animal Care.

### Extracellular recording

Recordings were performed using 32-channel silcon probes (NeuroNexus), in most cases polytrodes (A1x32-Poly2-5mm-50s-177) or occasionally linear arrays (A1x32-6mm-100-177), advanced with a stepping motor microdrive (M. Walsh Electronics, uD-800A). Raw electrophysiological signals were acquired with a Plexon Recorder (3 Hz to 8 kHz; sampling rate, 40 kHz), along with supplementary signals from a small photocell placed over one corner of the visual stimulus CRT, which were used for temporal registration of stimuli and spikes, and to verify the absence of dropped frames. Spike waveforms were carefully classified from the recorded multichannel data into single units, using Spikesorter (Swindale & Spacek, 2014). Only clearly sorted units were used for further analysis.

In total, 110 single units from 37 penetrations in 8 cats (4 males, 4 females) were analyzed. These recording experiments involved lab personnel working on other projects. Out of these neurons, 6 were rejected because part of their receptive fields was outside the screen, and 30 were rejected because the predictive performance of the fitted model was too low (see the Model Architecture section, below). The sample size included the remaining 74 neurons.

### Visual stimuli

Visual stimuli were presented on a gamma-corrected CRT monitor (NEC FP1350, 20 inches, 640x480 pixels, 150 Hz, 36 cd/m^2^) at a viewing distance of 57 cm. Stimuli were produced by an Apple Macintosh computer (MacPro, 2.66 GHz, 6 GB, MacOSX ver. 10.6.8, NVIDIA GeForce GT 120) using custom software written in MATLAB (ver. 2012b) with the Psychophysics Toolbox (ver. 3.0.10; Pelli, 1997; Brainard, 1997; Kleiner et al., 2007). We selected a channel having with good spike responses to hand-held bar stimuli, which we used to determine the dominant eye (with the non-dominant eye subsequently occluded), and to position the CRT monitor to be approximately centered around the population receptive field.

Visual stimuli were ensembles of 375 natural images taken from the McGill Calibrated Colour Image Database (Olmos & Kingdom, 2004), cropped to 480x480, converted to monochrome 8-bit integers - as in Talebi & Baker (2012), but with a higher RMS contrast. We randomly presented each ensemble at 75 images per second (i.e. every 13.33 ms) in short movies of 5 seconds each. We separated the ensembles into three sets, to evaluate predictive performance independently from overfitting. The training set had 20 movies which were presented 5 times each, while the validation and testing sets each had 5 movies which were presented 20 times each. The validation and testing sets were presented more often to provide less noisy estimates of the fitted model’s predictive performance. Instances of these three subsets of movies were quasi-randomly interleaved throughout the 45-minute recording session.

For the subsequent data analysis (described below), all images were resized from 480x480 to 40x40 before training (see below) to avoid overparameterization of the fitted model. Resizing was done using the Image module from the Python Image Library (PIL; Umesh, 2012).

### Model architecture

To better understand differences between the ON and OFF pathways, we employ a model architecture abstracted from known visual circuitry (Figure 1), whose parameters are optimized to predict a recorded cortical neuron’s mean spiking responses to the natural image ensembles. We model LGN receptive fields as parametrized 2D isotropic gaussians, acting convolutionally on the stimulus images. The antagonistic surrounds are neglected, so there is only a pair of gaussian width parameters, for the ON and OFF pathways, to be estimated. The connections between the gaussian operators and the model cortical neuron are a pair of linear weighted sums of rectified responses of the gaussian-operators, across a series of time lags. Each of these linear weighted sums acts like a “dense layer” in machine learning - but note that there is *not* a subsequent rectification. The output of each dense layer might be thought of as a presynaptic membrane potential contribution, from its respective ON or OFF pathway.

**Figure 1.**
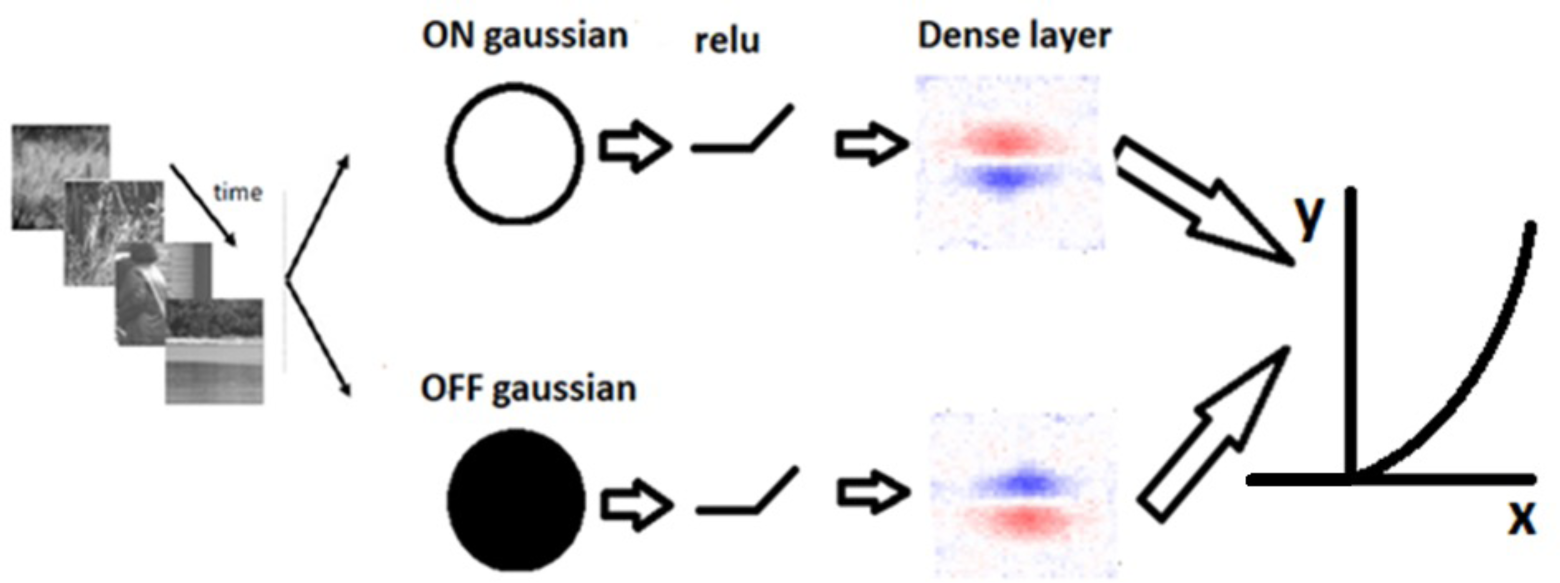
Model architecture for responses of a cortical neuron to visual stimuli such as natural images. Light and dark image regions are encoded as rectified responses of convolution with positive and negative spatial gaussians, respectively. Linear weighted sums are separately taken for each pathway and summed, followed by a half-power pointwise nonlinearity. A machine learning algorithm estimates the sizes of the parameterized gaussian operators, and the two sets of dense layer weights, for each of a series of time lags.

The inputs to the model are the pixel luminance values of the natural image stimuli (cropped, and spatially downsampled to 40x40 pixels, as described below). The mean of the inputs is centered at zero by subtracting the overall mean across all images within an ensemble. To model the neuron’s temporal processing, the inputs to the estimated model (see below) are composed of the preceding 7 images, each of which were presented for 13.33 ms. The model output is the neuron’s response, with spike times collected into time bins of 13.33 ms each (duration of each stimulus image frame).

The stimulus images are convolved with a pair of parametrized 2D gaussian filters (with positive or negative polarity for the ON and OFF pathways, respectively), each followed with a half-wave rectification (ReLU). The 2D gaussians represent receptive fields of LGN neurons in which the weaker surrounds (Croner & Kaplan, 1995) are neglected, as follows:

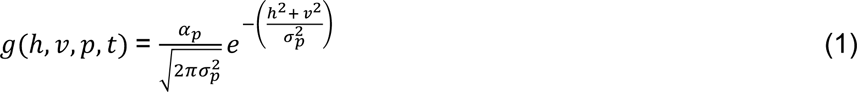

where h and v are the horizontal and vertical distances between a pixel and the center of the gaussian, respectively, and σ represents the standard deviation (i.e. width) and α the amplitude (i.e. the height) of the 2D gaussian. The σ parameter is estimated separately for each pathway (*p*). To allow each pathway to selectively process light or dark information, α is set to 1 for the ON pathway and -1 for the OFF pathway. The convolution of the 2D gaussian with the inputs is half-wave rectified (ReLU), to mimic spike frequency responses of LGN neurons (Persi et al., 2011):

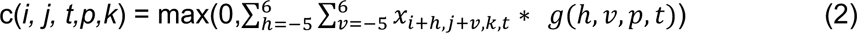

where *i* and *j* are the horizontal and vertical coordinates of the center of the 12x12 2D gaussian, *x* is the luminance of a specific pixel (resized to a grid of 40x40), *t* is the number of time bins between the shown image and the recorded response (latency), and *k* is the time bin of the neuron’s response. The convolution with the 2D gaussians is implemented with zero-padding and a “stride” of 1. Due to the first rectification, the ON-pathway encodes luminance above (lighter than) the mean, and the OFF-pathway luminance below (darker than) the mean.

For each of the ON- or OFF-pathways, the model then takes a linear weighted sum of the convolution outputs from the respective rectified gaussians, with each weight notionally representing the excitatory or inhibitory inputs from an array of LGN cells to the cortical neuron. The sum of responses from these dense layers is followed by a rectified power law output nonlinearity, which forms the final output of the model and the prediction of the neuron’s mean spiking response:

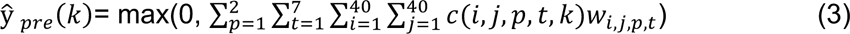

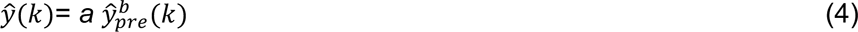

where *w* represents the dense layer weights, ŷ(*k*) is the prediction of a neuron’s response for the *k*^th^ time bin, *b* is the exponent of the rectified power nonlinearity, and *a* is a scale (gain) factor.

To estimate the proportion of the variance in a neuron’s response that is accounted for by the model’s predictions, we calculate a variance-accounted-for (VAF) index by taking the square of the Pearson correlation coefficient between y (neuron’s response) and ŷ (model’s predictions). To insure that the estimated weights are representative of each neuron’s responses to visual stimuli, we excluded neurons with a VAF below 10% in the testing set (see below). Based on this criterion we excluded 30 neurons, which resulted in a sample size of 74 neurons for the remaining analysis.

### Optimization and regularization

To characterize a neuron’s receptive field, we find the model parameters which minimize the difference between its recorded responses and the responses predicted by the model of Equation 3, which requires fitting a total of 2x40x40x7 = 22,400 dense layer weights, and 2 parameters (σ_*ON*_ *and* σ_*OFF*_) for the 2D gaussians (fitting of the two parameters for the output nonlinearity, Equation 4, is described below). To minimize over-fitting due to the large number of parameters, we employ L2-regularization by penalizing the squared amplitude of the dense layer weights (Hoel & Kennard, 1970), implemented by minimizing a loss function in which the first term is the squared error of the model prediction and the second term the regularization penalty:

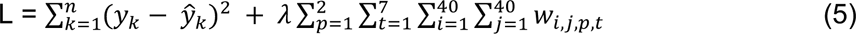

where ŷ_*k*_ is the neuron’s recorded response, ŷ_*k*_ the model’s predicted response for the *k*^th^ time bin, *w_i,j,t,p_* the dense layer weights (for the *p*-th On/Off stream, *t*-th time lag, and *i,j*-th dense layer position) and λ the L2-regularization hyperparameter. Based on pilot results from a representative subset of neurons, the hyperparameter λ is set to 5 x 10^-6^ in a first pass and 2 x 10^-6^ in a second pass (see below, three-pass training procedure). In the third pass, we train the model with different λ values of [1,2,4,8,16] x 10^-6^, and choose the λ value giving the best model performance on the validation dataset for each neuron.

This loss function is minimized using the Adam optimization algorithm (Kingma & Ba, 2014) with mini-batch gradient descent (Li et al., 2014). To further reduce overfitting, we apply dropout during training to both the convolutional and dense layers with a probability of 50% (Srivasta et al., 2014).

The data is separated into training, validation, and test sets, corresponding to the three sets of stimulus movies. The model parameters are fit to the training set using a mini-batch size of 100 stimulus-response pairs. As an additional regularization measure, training is stopped if there is no improvement on the validation set in the preceding 50 epochs - then we use the model at its peak performance (i.e. 50 epochs before training stops) in subsequent analyses. We use a third, separate test set to obtain an unbiased estimate of predictive performance.

### Three-pass training procedure

Because V1 receptive fields usually only occupy a small subset of the displayed visual stimulus images, it would be detrimental to optimize each neuron’s model based on the full extent of the images. Doing so would entail a very high overparameterization, or a loss of spatial resolution due to excessive downsampling of the stimulus images, in either case yielding poorer predictive performance. To address this issue, we use a three-pass training procedure, with each pass improving the spatial resolution of the receptive field estimate. In the first pass, we optimize the model parameters using the full 480x480 stimulus images downsampled to 40x40. We then manually designate a square cropping window that encloses an area slightly larger than the apparent receptive field. Next, we crop the stimulus images within that window, and rescale each image within it to 40x40. Due to the resizing, this cropped image then has much better spatial resolution than the 40x40 image from the first-pass. This image is used to re-train the model in the second pass, where we repeat the procedure, but with the cropped image. In the third pass, we further adjust the cropping window based on the model estimate obtained in the second pass. This third pass provides much higher accuracy in identifying the boundaries of the receptive field, and gives us the final model fits that we use for the remaining analysis. This three-pass training procedure allows us to characterize a neuron’s receptive field with high resolution and substantially increases predictive performance.

### Output nonlinearity

A cortical neuron’s spike frequency response has often been modeled with a final output nonlinearity, consisting of a rectified power law (Heeger, 1991; Anzai et al, 1999; Persi et al., 2011). However, it has proven problematic to simultaneously estimate the power law exponent with the other parameters using the backpropagation algorithm employed here. This problem is most likely due in part to the “exploding gradient” problem (Pascanu, Mikolov & Bengio, 2012). To resolve this issue, we initially set a power-law exponent value of unity (1.0), and wait 100 epochs into the training algorithm, to get a rough estimate of the other parameter values. We then pause the model optimization, to fit the two parameters of the output nonlinearity to the predicted vs. measured neuron responses - and then resume full model parameter optimization, keeping the output nonlinearity parameters fixed.

To address the heavily uneven distribution of the measured firing rates, we bin the predicted responses into 100 bins of 75 responses each, and compute the mean measured response for each bin - a modification of the method used by Anzai et al (1999). We then fit a scaling factor ‘*a*’ and an exponent ‘*b*’ (Eq. 4) to minimize the difference between the binned predicted responses ŷ and the measured spike rates *y*, using python scipy’s ‘*optimize.curve_fit’*.

### Estimating excitation and inhibition

The spatiotemporal properties of each of the ON and OFF pathways in the fitted model depend on both the dense layer weights and estimated 2D gaussians (which may differ in width for the ON and OFF pathways). To incorporate both in our analysis, for each of the ON and OFF pathways we convolve the 2D gaussian with the corresponding dense weights, to produce a 40x40x7 spatio-temporal filter for each pathway (ON_Recon_ and OFF_Recon_). This “reconstructed” receptive field represents the neuron’s responsiveness to either light or dark stimuli. For further analyses we estimate the overall amount of excitation and inhibition from the filter for each pathway and time lag, by taking the sum of all positive or negative values in either the ON or OFF reconstructed receptive field. This procedure provides an inference of the total amount of ON excitation, ON inhibition, OFF excitation and OFF inhibition contributing to each neuron’s response:

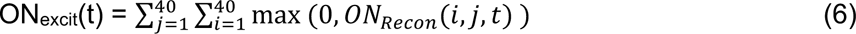

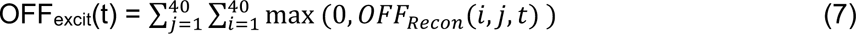

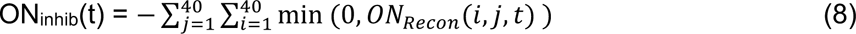

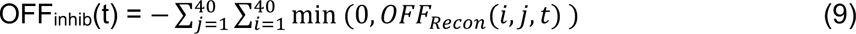

where ON_Recon_ represents the contribution of the ON pathway to the estimated relationship between (light) stimuli and the neuron’s responses (and similarly for OFF_Recon_). ON_Recon_ and OFF_Recon_ are estimated from the convolution of the spatiotemporal weights, *w_p,t_*, with the gaussian layer, *g_p,t_* for *p*=1 (and similarly for OFF_Recon_ for *p*=2):

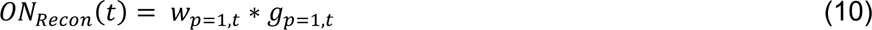

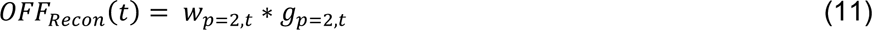

Parts of our analysis are based on each neuron’s peak latency of responsiveness, determined as the latency having the greatest variance in the sum of the reconstructions from each pathway:

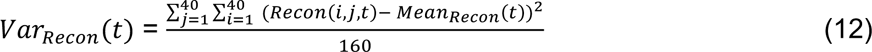

where

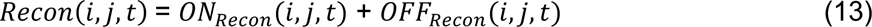

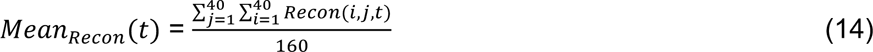

The latency *t* with the highest *VAR_Recon_* will be referred to as T.

### Light/dark balance

To quantify the extent to which individual neurons are light- or dark-dominated, we use a light-dark balance index (LDB) to indicate the relative influence of a neuron’s light and dark weights:

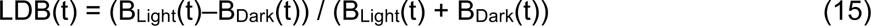

where

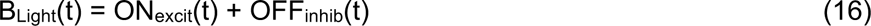

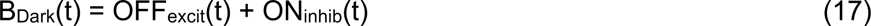

This index varies from -1.0 to 1.0, with positive LDB values indicating a neuron is light-dominated, and negative values that it is dark-dominated.

### Excitation/Inhibition balance

The excitation/inhibition balance (EIB) index is similar to LDB, but contrasts excitation with inhibition instead of light with dark:

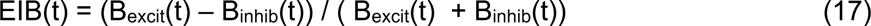

where

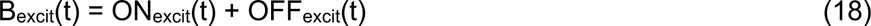

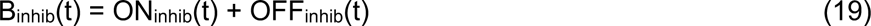

This index varies from -1 to 1, with positive EIB values indicating a neuron’s response reflects relatively stronger excitation, and negative EIB values stronger inhibition.

### Simulated responses to artificial stimuli

To better understand how dark-dominance influences neurons’ responses to visual stimuli, we simulated the estimated models’ responses to four different stimulus conditions. The 40x40 stimuli were tailored to each neuron’s spatial receptive field, which we estimated by using Recon(i,j,T) (Eq. 13) at each neuron’s peak latency T (see above). The four stimulus conditions (Fig. 6) are the following: light falling on light-driven regions (LL), dark on dark-driven regions (DD), light on light-driven and half of dark-driven regions (LLHD) and dark on dark-driven and half of light-driven regions (DDHL):

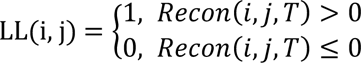

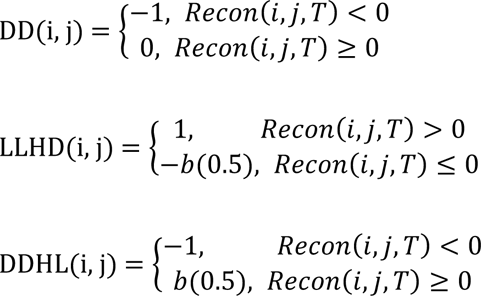

where i and j index the spatial location of the image pixels, T is each neuron’s peak latency and b is a random value (either 0 or 1) drawn from an equiprobable Bernoulli distribution. The first two stimulus conditions (LL and DD) are designed to only recruit excitation, while the latter two conditions recruit a mixture of both excitation and half as much inhibition. We used the estimated model of each neuron to simulate its response to each of these four stimulus conditions at different latencies. (Note that this procedure is equivalent to simulating the neuron’s impulse response to each stimulus.) The average simulated responses across the entire sample at each latency are shown in Figure 6.

### Orientation selectivity

To better understand the relationship between dark-dominance and orientation selectivity, we simulated the estimated models’ responses to static sinewave gratings at each of 36 orientations (with increments of 5 degrees), 56 spatial frequencies (equally spaced from 0.0667 to 0.143 cycles per image) and 36 phases (increments of 5 degrees). These responses were used to compute the orientation selectivity of each neuron using a vector summation method (Wörgötter & Eysel, 1987; Swindale, 1998):

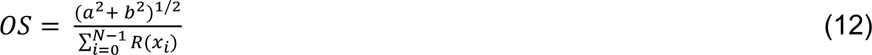

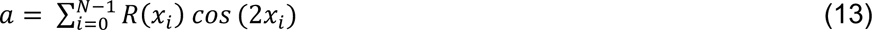

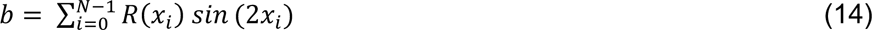

where N is the number of sinewave gratings, xi is the orientation angle, and R(xi) represents the simulated responses. The orientation selectivity index, *OS*, was computed separately for each latency in individual neurons.

### Experimental design and statistical analyses

Most statistical tests here are paired t-tests, to compare whether there is a significant difference between the means of two groups. We also use one-sample t-tests to assess whether means differ significantly from zero, and perform linear regression to test the correlation between two sets of values. We adjust for multiple comparisons with Bonferroni corrections, where the significance threshold α of 0.05 is divided by the number of comparisons (e.g., Figure 3B has 6 comparisons: α = 0.05 / 6 = 0.0083). Because visual responses are much weaker for latencies longer than 40 ms (see Figure 2), statistical tests are only performed for the first three latencies in Figures 5 and 7, with the correction for multiple comparisons adjusted accordingly.

**Figure 2.**
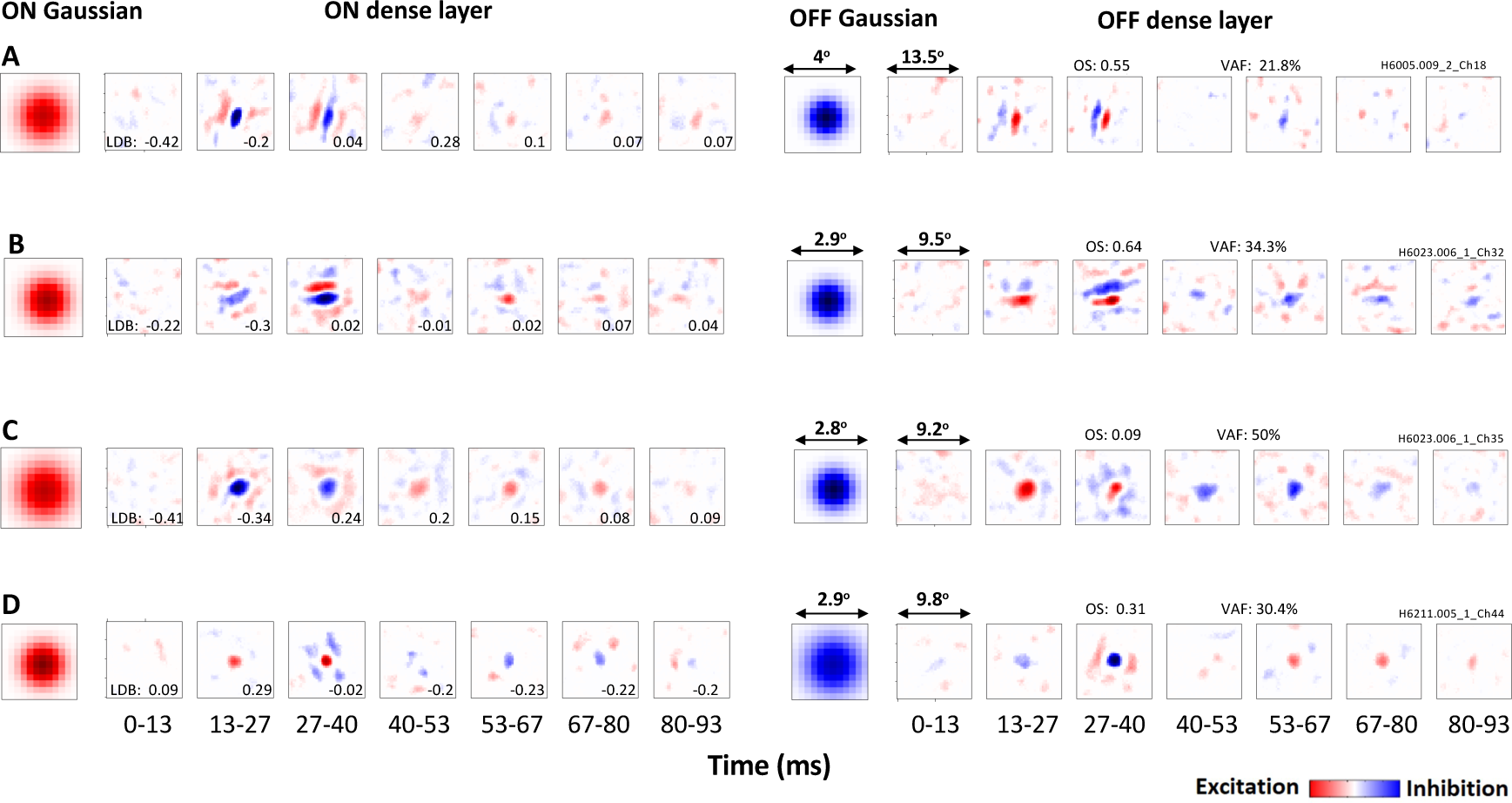
Gaussian filters and spatiotemporal dense layers estimated for four example neurons, one in each row. Elements of ON pathway on the left and OFF pathway on the right. Each dense layer is shown at a series of latencies ranging from 0-13 ms to 80-93 ms, with neurons being most responsive at the 13-27 ms and 27-40 ms latencies. Positive values (excitation) are in red and negative values (inhibition) are in blue. Orientation selectivity (OS) and variance accounted for (VAF) are indicated for each neuron, and light-dark balance (LDB) values for each latency. ***A, B***, Both neurons respond more strongly to dark stimuli (LDB < 0) at the 13-27 ms latency, and become more balanced (LDB ∼ zero) at the 27-40 ms latency. ***C***, Neuron is also dark-dominant at the 13-27 ms latency but responds more strongly to light at the 27-40 ms latency. ***D***, Neuron that is instead light-dominant at the 13-27 ms latency, and balanced at the 27-40 ms latency.

**Figure 3.**
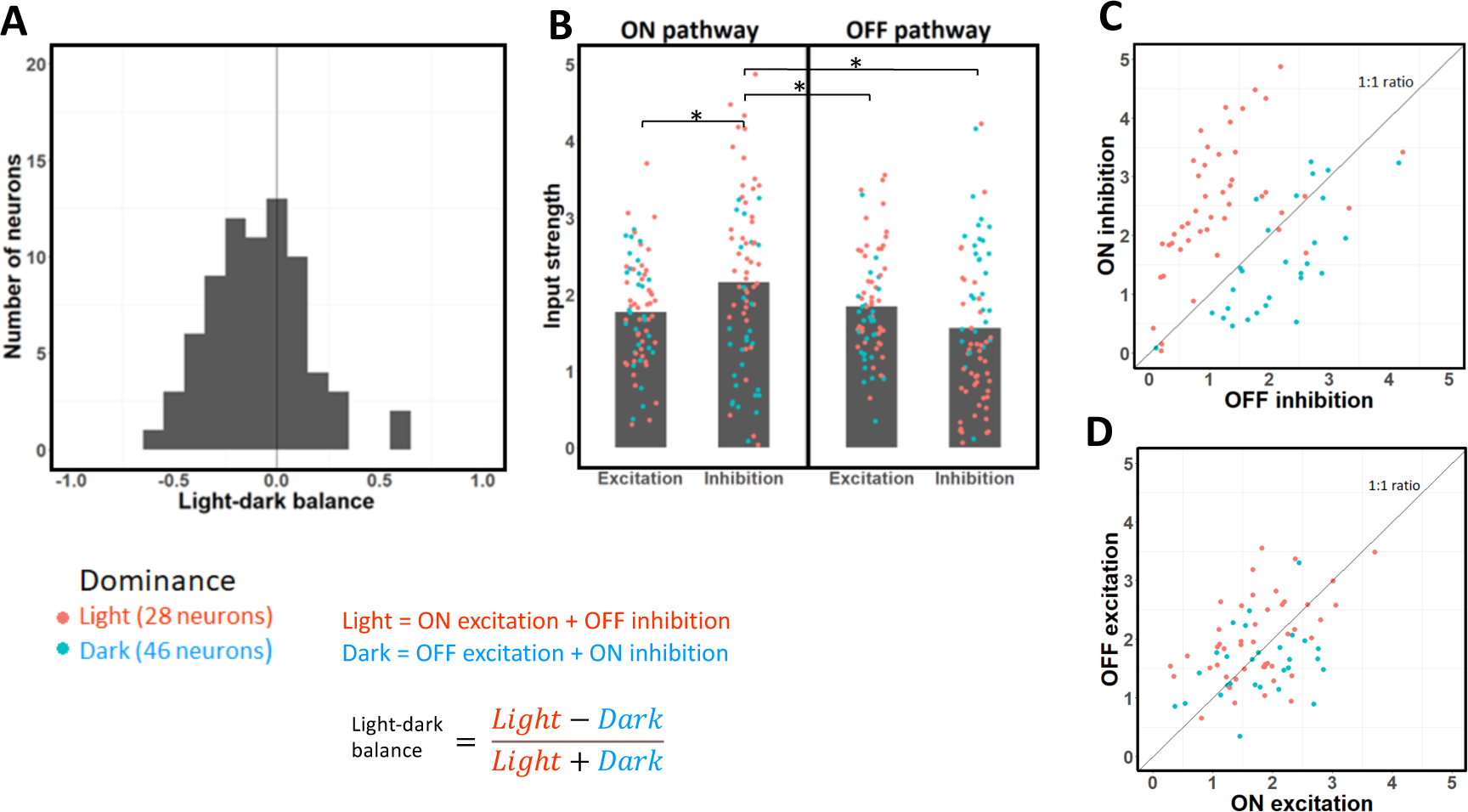
Strengths of excitation and inhibition from the ON and OFF pathways at each neuron’s optimal time lag. ***A***, Distribution of light-dark balance (LDB) index values for each neuron at its optimal latency. This index is on average negative, which indicates neurons respond more strongly to dark than light stimuli. ***B***, Strength of excitation and inhibition across the ON and OFF pathways for each neuron, with average values shown as gray bars. Note ON inhibition is the strongest input on average. Blue dots represent light-dominant neurons and orange dots represent dark-dominant neurons. Significant paired t-tests (with Bonferroni correction, p < 0.0083) are indicated by a star (*). ***C***, Scatterplot of ON vs. OFF inhibition, for each of the 74 neurons. Most neurons have stronger ON inhibition, and whether ON or OFF inhibition is stronger is correlated with light and dark-dominance. ***D***, same as (*C*) but for ON and OFF excitation. Unlike the result for inhibition in (*C*), note that ON and OFF excitation have relatively similar strength on average.

## Results

As described in the Methods, a simple neural network model (Figure 1) was fit to responses from individual neurons, to estimate 2D gaussians and 3D spatiotemporal filters (dense layers) separately for ON and OFF inputs, as well as a power law output nonlinearity. Figure 2 shows these estimated model parameters for four example neurons, which all had peak responses at the 13-27 or 27-40 ms latency. As we observed more generally, the early ON and OFF Gaussian filters for a given neuron were about the same size, but opposite in polarity. And for each neuron, the spatiotemporal filters (dense layers) were largely similar, both spatially and temporally, but opposite in polarity.

Many of the neurons had Gabor-like receptive fields that are orientation selective (Hubel & Wiesel, 1962), like the one shown in Figure 2A (with OS = 0.55). At the 27-40 ms latency, this neuron has balanced light and dark responses for both the ON (left) and OFF (right) pathways (LDB = 0.04). This balance does not occur at the 13-27 ms latency, where the neuron responds more strongly to dark stimuli (LDB = -0.2). This bias is due to the OFF pathway having stronger excitation (red) than inhibition (blue), with the ON pathway being balanced.

Another neuron (Figure 2B) is also orientation selective (OS = 0.64), balanced (LDB = 0.02) at the 27-40 ms latency, and exhibits a bias toward dark responses (LDB = -0.3) at the 13-27 ms latency. However, for this neuron the 27-40 ms latency is imbalanced due to both the ON and OFF pathways, with the ON pathway having weaker excitation and the OFF pathway having weaker inhibition.

The neuron shown in Figure 2C differs from the previous examples in that it has low orientation selectivity (OS = 0.09) due to its isotropic receptive field, which has a dark center and an opposite-polarity surround. At the 13-27 ms latency, this neuron is dark-dominant (LDB = -0.34) due to its weaker surround, especially in the OFF pathway. Contrary to the above two example neurons, at the 27-40 ms latency this neuron is not balanced but light-dominant, due to stronger inhibition than excitation in the OFF pathway.

Not all neurons are dark-dominant - for example, the neuron in Figure 2D is light-dominant at the 13-27 ms latency (LDB = 0.29), due to its Gaussian-like receptive field with a light-responsive center and a weak surround. Similar to previous results, this neuron is balanced at the 27-40 ms latency (LDB = -0.02). However, as we shall see below, there is a tendency for most neurons to, on average, have stronger responses to dark stimuli at the 13-27 ms latency, and to have stronger responses to light stimuli or to be balanced at the 27-40 ms latency.

### Population responses

To investigate the patterns of light and dark response strength across the sample of 74 neurons, we computed the sums of the four types of inputs for each neuron’s optimal time lag (see Methods). Neurons had an optimal time latency of either 0-13.3 (38 neurons), 13.3-26.7 ms (35 neurons) or 26.7-40 ms (1 neuron). As described in the Methods, we estimated the overall amount of excitation and inhibition from the ON and OFF pathways, and also used these values to calculate an index of light-vs-dark balance, LDB. We classified each neuron as dark-dominated (LDB < 0) or light-dominated (LDB > 0) depending on whether it was more responsive to dark (OFF excitation and ON inhibition) or light (ON excitation and OFF inhibition) at its optimal time latency. Across our population of 74 neurons, we found 46 neurons (62.16%) to be dark-dominated (LDB<0) and 28 neurons (37.84%) to be light-dominated (LDB>0) at their optimal latencies, similar to Yeh et al. (2009). The neurons in our sample had a wide range of LDB values (Figure 3A; minimum = -0.62, maximum = 0.57, median = -0.078), but were on average dark-dominated, with an average LDB of -0.094 (Figure 3A; t = -3.49, df = 73, p = 0.00081).

To better understand why cortical neurons are on average more responsive to dark than light stimuli, we next compare the four types of inputs (Figure 3B) at each neuron’s optimal latency. ON pathway inhibition is the strongest type of input on average, and is significantly stronger than the other three. ON inhibition is on average significantly stronger than ON excitation (paired t-tests with Bonferroni correction; t = 4.21, df = 73, p = 7.07 x 10^-5^), OFF excitation (t = 3.03, df = 73, p = 0.0034) and OFF inhibition (t = 3.95, df = 73, p = 0.00018). In contrast, OFF inhibition is on average the weakest type of input. While it has previously been suggested that stronger OFF than ON excitation could underlie stronger dark responses (Jin et al., 2008), the overall dark-dominance effect we observe at the optimal latency instead seems to be due to a strong imbalance between ON and OFF inhibition: while inhibition is on average 37.95% stronger from the ON than from the OFF pathway (Figure 3C), there is no significant difference between excitation from the ON and OFF pathways (Figure 3D; t = 0.81, df = 73, p = 0.42). The difference between ON and OFF inhibition (Figure 3C) is also significantly stronger (t = 2.96, df = 73, p = 0.0042) than the difference between ON and OFF excitation (Figure 3D).

In addition, whether a given neuron is light- or dark-dominated is strongly related to whether ON inhibition exceeds OFF inhibition (Figure 3C, red points above 1:1 line vs. blue points below). However, the imbalance of ON vs. OFF excitation poorly predicts whether a neuron is light or dark-dominant (Figure 3D). Overall, these results suggest the dark-dominance effect to be more driven by an imbalance in ON/OFF inhibition than by an imbalance in ON/OFF excitation.

### Time dynamics

Since responses to dark stimuli have previously been found to have shorter latencies than responses to light stimuli (Komban et al., 2014), we suspected the above results might vary as a function of response latency. The dependence of light-dark balance is shown for each of the measured time lags in Figure 4A, with data points for each sampled neuron, and gray bars indicating their averages. The dark-dominance effect is especially predominant at the 0-13.3 ms (one sample t-tests with Bonferroni correction; t = -7.68, df = 73, p = 5.6 x 10^-11^) and 13.3-26.7 ms latencies (t = -3.90, df = 73, p-value = 0.00021). The dark-dominance effect disappears at the 26.7-40 ms latency, with slightly stronger average responses to light than dark, though the difference is not significant (t = 2.48, df = 73, p = 0.015). At the longer latencies, there is no significant average light- or dark-dominance (p > 0.15). These findings suggest that while V1 neurons are on average biased towards dark responses in their short latencies, the dark-dominance effect disappears at the 27-40 ms latency.

**Figure 4.**
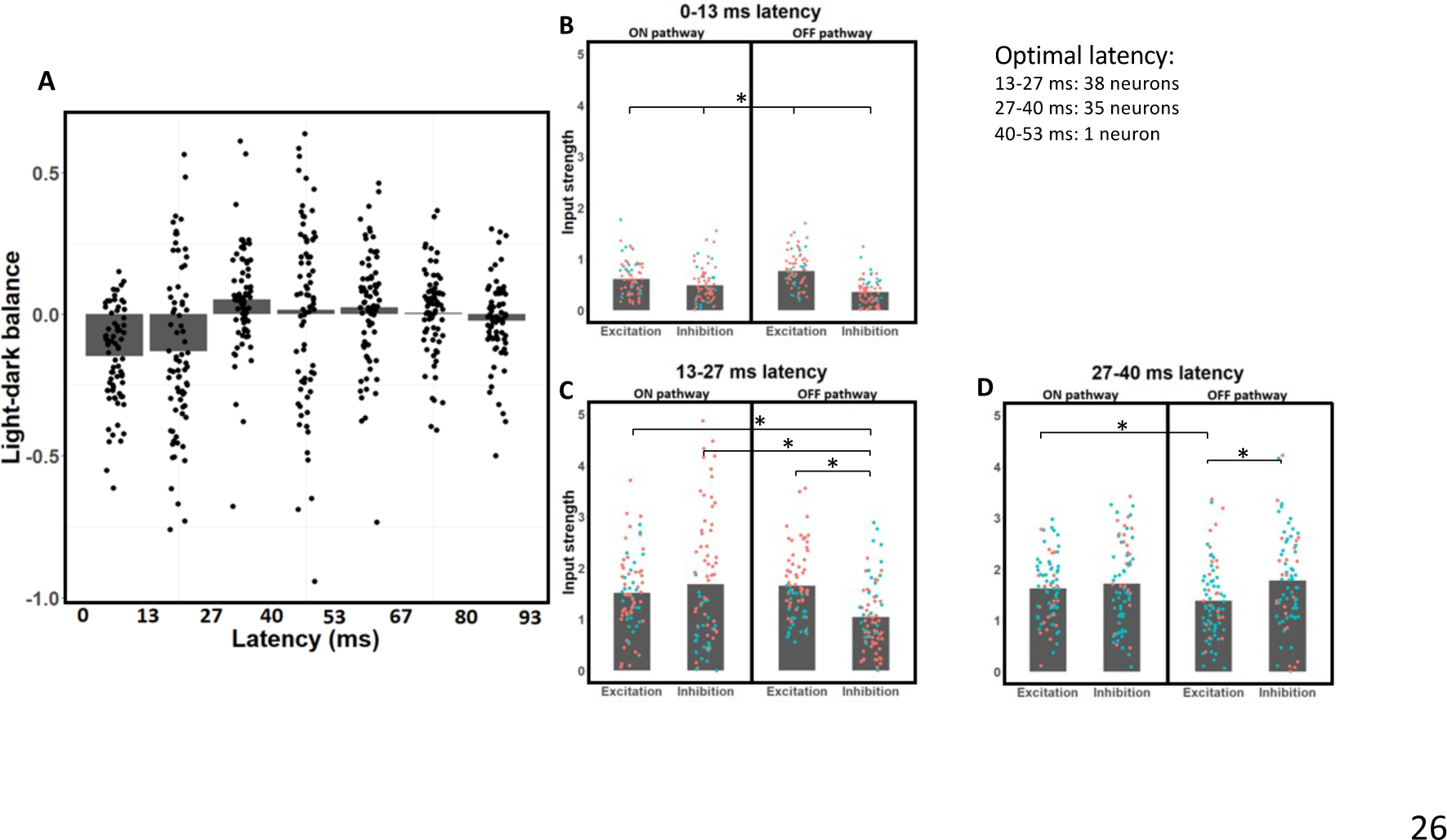
Light-dark balance index and strength of excitation/inhibition of ON and OFF pathways for all neurons, across different latencies. ***A***, Light-dark balance index values, shown as bar graph of average values for each latency, with superimposed data points for individual neurons. The 0-13.3 and 13.3-26.7 ms latencies exhibit dark-dominance, the 26.7-40 ms latency shows a slight bias towards light-dominance and the later latencies are relatively balanced. ***B***, Excitation and inhibition from the ON and OFF pathways for each neuron at the 0-13.3 ms latency. OFF excitation is stronger than ON excitation on average, and inhibition is significantly weaker than excitation at this latency. ***C***, Same as (*B*) but for the 13.3-26.7 ms latency - note the relatively balanced values on average, except for OFF inhibition which is significantly weaker than the other three types of input. ***D***, Same as (*B,C*) but for the 26.7-40 ms latency - note the significantly weaker OFF excitation on average compared to the other three types of input. Significant paired t-tests (with Bonferroni correction, p < 0.0083) are shown by a star (*) for *B, C* and *D*. All of the pair-wise comparisons are significantly different in *D*.

To understand why neurons are dark-dominated in their early latencies, we investigate how the strength of each input type varies as a function of time. Because neurons are most responsive up until a latency of 40 ms, the following sections focus on the first three latencies. As we can see in Figure 4B, at a latency of 0-13.3 ms OFF excitation is the strongest input on average - it is 25.5% stronger than ON excitation (paired t-tests with Bonferroni correction; t = 5.47, df = 73, p = 6.0 x 10^-7^). Inhibition is significantly weaker than excitation, both in the ON (t = 4.35, df = 73, p = 4.36 x 10^-5^) and OFF (t = 13.1, df = 73, p < 2.2 x 10^-16^) pathways. This discrepancy is stronger in the OFF than in the ON pathway (t = 7.51, df = 73, p = 1.17 x 10^-10^). Inhibition is on average 37.9% stronger from the ON than from the OFF pathway (t = 5.35, df = 73, p = 9.76 x 10^-7^), thereby contributing to stronger dark responses. Weaker inhibition than excitation at the shortest latency could be explained by inhibition having to go through at least one more synapse than excitation to reach V1 neurons (Ferster & Lindström 1983, Martin & Whitteridge 1984; Montero, 1986). These results are also consistent with findings from Jin et al. (2008), who demonstrated stronger OFF than ON excitation from the LGN to be an important mechanism contributing to the dark-dominance phenomenon. However, while stronger OFF than ON excitation might explain dark-dominance at the 0-13.3 ms latency, the overall dark/light dominance of neurons will be more related to the considerably stronger responses at the 13.3-26.7 and 26.7-40 ms latencies.

Responses at the 13.3-26.7 ms latency are also stronger to dark stimuli (Figure 4C), but for a different reason. OFF excitation is not significantly stronger than ON excitation at the 13.3-26.7 ms latency (paired t-tests with Bonferroni correction; t = 1.65, df = 73, p = 0.103). Instead, dark-dominance at this latency is due to weaker OFF inhibition compared to the other three types of inputs. Inhibition from the OFF pathway is on average 38.3% weaker than inhibition from the ON pathway (t = 4.58, df = 73, p = 1.88 x 10^-5^). OFF inhibition is also on average 31.2% weaker than ON excitation (t = 7.67, df = 73, p = 5.77 x 10^-11^) and on average 36.9% weaker than OFF excitation (t = 5.84, df = 73, p = 1.37 x 10^-7^). No other pair of inputs are significantly different from each other (p < 0.0083) at the 13.3-26.7 ms latency, further strengthening the idea that the imbalance between light and dark responses at this latency is due to weaker OFF inhibition.

Contrary to the results for the earlier latencies, the 26.7-40 ms latency does not show dark-dominance (Figure 4D). The only significant differences are OFF excitation being both 22.4% weaker than OFF inhibition (paired t-tests with Bonferroni correction; t = 5.37, df = 73, p = 9.1 x 10^-7^) and 19.7% weaker than ON inhibition (t = 3.80, df = 73, p = 0.0003). No other pair of inputs differ significantly (p < 0.0083) at this latency. Thus, different types of input are relatively more balanced at this latency compared to the previous ones.

To further understand the time dynamics of dark and light responses, we next analyze how the strength of each type of input changes across latencies (Figure 5). Because all input types are much weaker at the 0-13.3 ms latency compared to the 13.3-26.7 and 26.7-40 ms latencies (Figure 5), we focus our analysis on comparing the two latencies with the strongest responses, 13.3-26.7 ms and 26.7-40 ms. Consistent with the above results, OFF inhibition is 41.4% weaker at 13.3-26.7 ms than at 26.7-40 ms (Figure 5A; paired t-tests with Bonferroni correction; t = 8.14, df = 73, p = 7.63 x 10^-12^), and is the only input type to differ significantly in strength between these two latencies. OFF excitation is 16.6% weaker at 26.7-40 ms than at 13.3-26.7 ms, but this difference is not significant (Figure 5B; t = 2.11, df = 73, p = 0.0381). There are no significant differences between the 13.3-26.7 ms and 26.7-40 ms latencies for both ON inhibition (Figure 5C; t = 0.197, df = 73, p = 0.845) and ON excitation (Figure 5D; t = 1.15, df = 73, p = 0.256). These findings suggest that inhibition is slower to dark than light stimuli, which leads to dark-dominance at the 13.3-26.7 ms latency.

**Figure 5.**
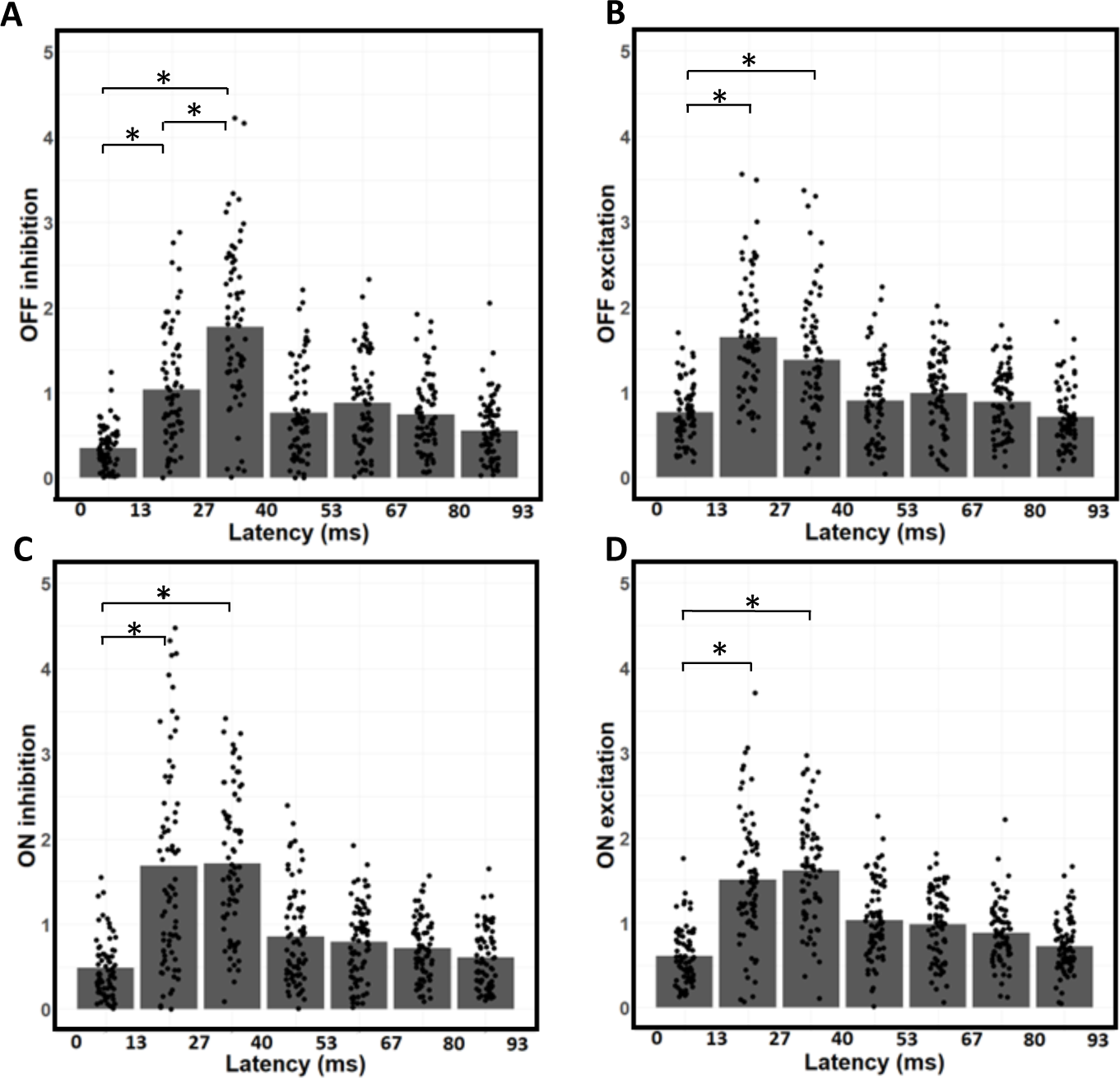
Temporal dependence of contributions from ON / OFF excitation and inhibition. ***A***, Bar graph of average OFF inhibition strength across time lags, with data points indicating values for individual neurons. Note OFF inhibition is weaker at the 13.3-26.7 ms latency than at the 26.7-40 ms latency. ***B***, Same as (*A*) but for OFF excitation. ***C***, Same as (*A-B*) but for ON inhibition. ***D***, Same as (*A-C*) but for OFF excitation. Significant paired t-tests (p < 0.0167) between the first three latencies are shown by a star (*). Note that OFF inhibition (*A*) is the only input type to significantly vary in strength between the 13.3-26.7 and 26.7-40 ms latencies. Also note that input strength at the 0-13.3 ms latency is always significantly weaker than at the 13.3-26.7 and 26.7-40 ms latencies.

**Figure 6.**
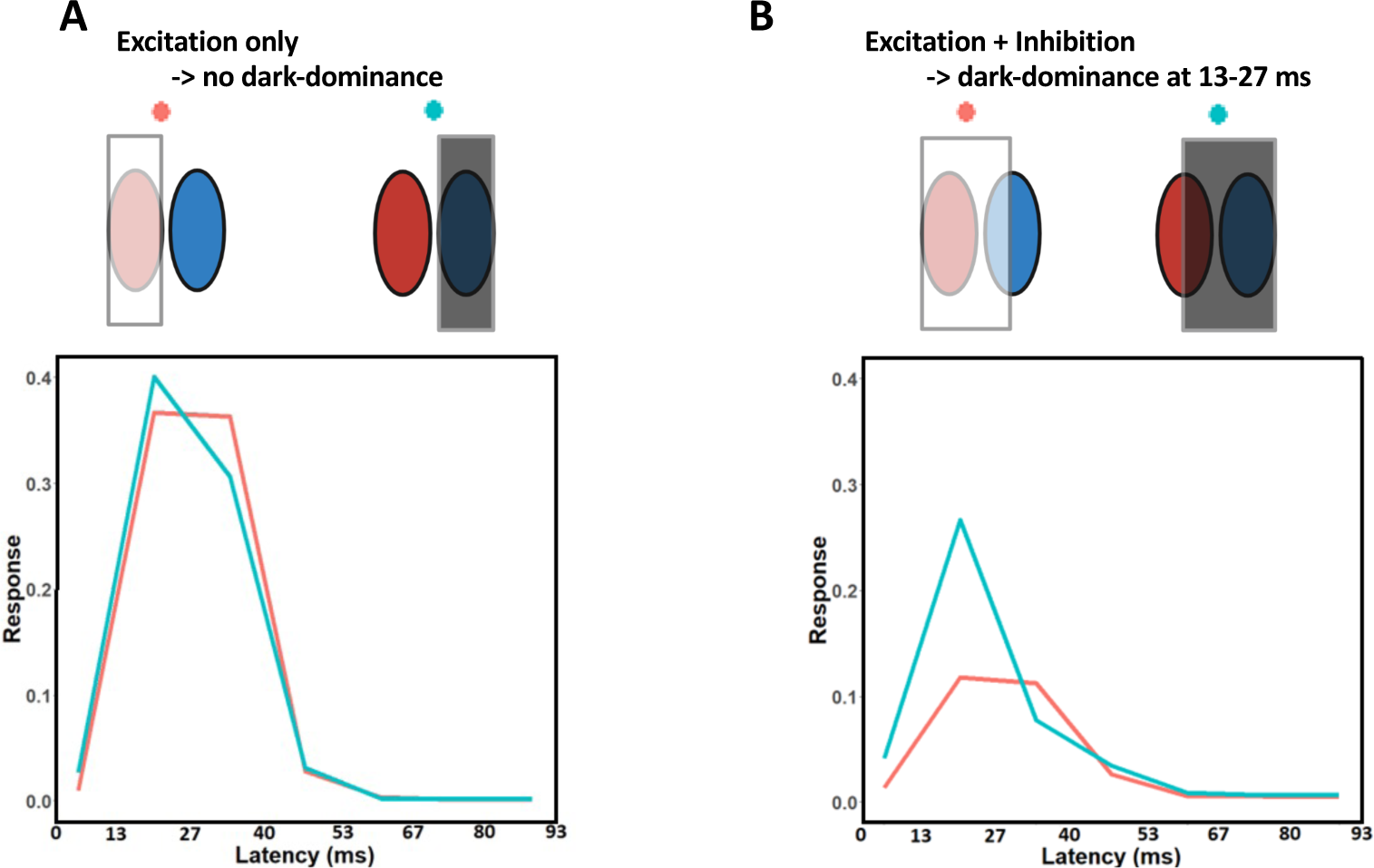
Average simulated temporal impulse responses to different stimuli. The stimuli were tailored to each neuron’s receptive field and are presented here in a schematic form. ***A***, Red shows the simulated responses to light stimuli on the light-driven regions of each neuron’s receptive field. Blue shows the simulated response to dark stimuli on the dark-driven regions of each receptive field. The two responses are similar, suggesting responses to light and dark stimuli are relatively balanced across latencies. ***B***, Red shows the simulated responses to light stimuli falling upon the light-driven regions, and also upon half of the dark-driven regions. Blue shows the simulated responses to dark stimuli on the dark-driven regions, plus on half of the light-driven regions. As expected, the responses are weaker than in (*A*), and this decrease is much less pronounced for the dark stimulus (blue line) at the 13-27 ms latency. These results suggest that dark-dominance predominantly occurs when measured with stimuli that recruit both the excitatory and inhibitory regions of a neuron’s receptive field.

**Figure 7.**
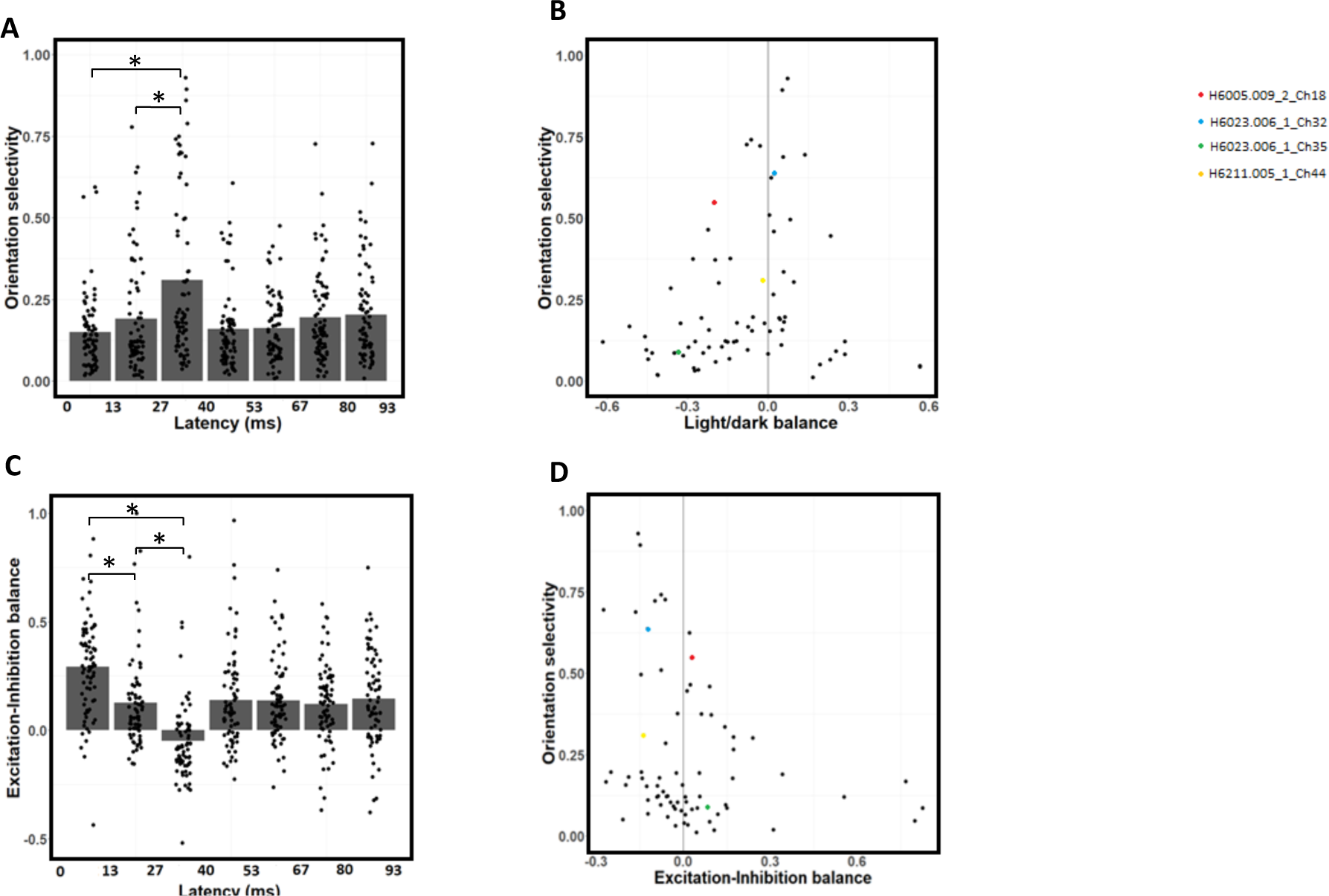
Changes in orientation selectivity and excitation-inhibition balance across latencies. ***A***, Average orientation selectivity peaks at the 26.7-40 ms latency, and is relatively low at the 0-13.3 and 13.3-26.7 ms latencies. ***B***, Relationship between orientation selectivity (ordinate) and light-dark balance (abscissa). Neurons with higher orientation selectivity tend to be more balanced. ***C***, Excitation-Inhibition balance (EIB) index as a function of latency. Excitation is stronger than inhibition at the 0-13.3 and 13.3-26.7 ms latencies, while excitation and inhibition are relatively balanced at the 26.7-40 ms latency. ***D***, Relationship between orientation selectivity (ordinate) and EIB (abscissa). Neurons with stronger excitation than inhibition tend to be less orientation selective. Significant paired t-tests (with Bonferroni correction, p < 0.0167) between the first three latencies (0-40 ms) are shown by a star (*) for *B* and *D*.

The results so far suggest dark-dominance occurs at the 13-27 ms latency due to more inhibition to light than dark stimuli. Because of those results, we hypothesized the dark-dominance effect would depend on how much inhibition a neuron receives, which in turn depends on the relationship between the stimuli and a neuron’s receptive field. There should be little or no dark-dominance from excitation alone, for example if we compare the responses to light stimuli falling upon the ON-excitation region to dark stimuli on the OFF-excitation region. Dark-dominance also cannot occur from inhibition alone, since the spontaneous firing rate is close to zero. Instead, dark-dominance should occur when a stimulus triggers both excitation and inhibition, for example when either light or dark stimuli fall upon both the light and dark-driven regions of a neuron’s receptive field.

To test this hypothesis, we simulated responses of the estimated models to four different stimulus conditions tailored to the receptive field of each neuron (see Methods): Light stimuli on light-driven regions, 2. Dark stimuli on dark-driven regions, 3. Light stimuli on light and (half of) dark-driven regions, and 4. Dark stimuli on dark and (half of) light-driven regions (Figure 6, top parts). The averages of the four responses were taken across the entire sample of 74 neurons (Figure 6, lower plots). As expected, the simulation shows little or no dark-dominance when only the excitatory region is stimulated (conditions 1 and 2, Figure 6A). But as we hypothesized, we do obtain dark-dominance at the 13-27 ms latency with stimuli that both excite and inhibit the neuron’s response (conditions 3 and 4, Figure 6B). These results support the idea that dark-dominance occurs when measured with stimuli that recruit both the excitatory and inhibitory regions of a neuron’s receptive field.

### Orientation selectivity

Previous studies demonstrated that V1 neurons are less orientation selective in their early responses (Ringach et al, 1997; Shapley et al, 2003). Since we have found early latencies to respond more strongly to dark stimuli, we wondered whether the stronger dark-dominance might be related to weaker orientation selectivity. To infer orientation selectivity, we next simulate the responses of the neurons’ fitted models to static sinewave grating stimuli with a series of orientations, spatial frequencies, and phases. For each latency, we select the sinewave grating with the best phase and spatial frequency for each orientation. We then use each model’s simulated responses to these sinewave gratings to measure an index of orientation selectivity, *OS*, using a conventional vector summation method (Wörgötter & Eysel, 1987; Swindale, 1998; see Methods), as a function of latency. This orientation selectivity index for a given neuron typically peaks at the 26.7-40 ms latency (Figure 7A). More specifically, across the population the orientation selectivity is significantly higher at 26.7-40 ms than at both 13.3-26.7 ms (paired t-test with Bonferroni correction; t = 6.07, df = 73, p = 5.1 x 10^-8^) and 0-13.3 ms (t = 5.95, df = 73, p = 8.4 x 10^-8^). Orientation selectivity is slightly higher at the 13.3-26.7 than at the 0-13.3 ms latency, but this difference is not significant (t = 2.17, df = 73, p = 0.033). These results suggest that orientation selectivity is most prominent at the 26.7-40 ms latency, where is also the first latency where light and dark responses are relatively balanced.

We next investigate the relationship between orientation selectivity and light-dark balance at each neuron’s optimal latency, which can be seen in Figure 7B. Neurons having high dark dominance (LDB << 0) or high light dominance (LDB >> 0) tend to have low orientation selectivity, while those that are more orientation selective are more often light-dark balanced (LDB ∼ 0). This apparent relationship is confirmed statistically: there is a significant negative relationship (r = -0.45) between orientation selectivity and absolute values of LDB (t = -4.4, df = 72, p = 4 x 10^-5^). These results suggest that a response bias towards dark stimuli might reduce a neuron’s orientation selectivity (Figure 7B), especially at the 0-13.3 and 13.3-26.7 ms latencies (Figure 7A).

Another possible explanation for weaker orientation selectivity at early latencies could be faster excitation than inhibition (Ringach et al., 1997; Shapley et al., 2003). Figure 7C shows the relative amount of excitation vs. inhibition (EIB index; see Methods) at each latency. Excitation is stronger than inhibition at the 0-13.3 ms (t = 11.3, df = 73, p < 2.2 x 10^-16^) and 13.3-26.7 ms latencies (t = 5.08, df = 73, p = 2.8 x 10^-6^), while there is no significant difference between excitation and inhibition at the 26.7-40 ms latency (t = -2.3, df = 73, p = 0.025). This bias towards excitation weakens over time, with lower EIB values for the 13.3-26.7 than for the 0-13.3 ms latency (t = 4.89, df = 73, p = 5.73 x 10^-6^). EIB values are also lower for the 26.7-40 than for the 13.3-26.7 ms latency (t = 7.44, df = 73, p = 1.58 x 10^-10^). Also consistent with Ringach et al. (1997) and Shapley et al. (2003), we find a negative correlation of r = -0.26 between orientation selectivity and EIB (Figure 7D; t = -4.87, df = 72, p = 6.25 x 10^-6^). Overall, these results suggest that both dark-dominance and stronger excitation contribute to weaker orientation selectivity at early latencies. However, another interpretation might be that weaker OFF inhibition is responsible for all of the above phenomena at early latencies: stronger dark responses, weaker overall inhibition and weaker orientation selectivity (see Discussion).

## Discussion

Using a novel model-fitting approach to natural image responses, we find V1 neurons respond more strongly to dark than to light stimuli at early but not at later latencies, due to slower inhibition to dark than light stimuli. Dark-dominance occurs when inhibition is differentially recruited, for example when there is a light stimulus on the dark-driven region of a neuron’s receptive field (or vice-versa). As can be seen in Figure 6 our results suggest little difference in the average neuron’s firing rate when a light stimulus only covers the light-excited region of the receptive field (Figure 6A, red) compared to when a dark stimulus only covers the dark-excited region of the receptive field (Figure 6A, blue). At the 13.3-26.7 ms latency, stronger responses to dark stimuli are instead observed when light (Figure 6B, red) or dark (Figure 6B, blue) stimuli cover both the light and dark regions of a neuron’s receptive field. These results could help explain why dark-dominance increases at lower spatial frequencies (Jansen et al., 2019), since a given light or dark band of a low-frequency grating may cover more than one region of a receptive field.

### Inference of excitation and inhibition from model-fitting

We use a machine learning algorithm to fit a model based on separate ON and OFF retinogeniculate inputs to V1, each composed of linear filters followed by half-wave rectification. The weaker surrounds of LGN neurons (e.g. Croner & Kaplan, 1995) are omitted, to enable robust convergence on a set of fitted parameter values. Using this approach, we can distinguish between excitation and inhibition to light and dark stimuli across spatial receptive field locations and temporal lags, to investigate how ON and OFF pathways contribute to the dark-dominance effect.

It is important to note that the excitation and inhibition we estimate does not necessarily reflect direct LGN inputs. For example, V1 does not receive direct inhibitory inputs from the LGN (Ferster & Lindström 1983, Martin & Whitteridge 1984; Montero, 1986), but rather from local inhibitory interneurons, which in turn may relay geniculate inputs or be driven by other V1 neurons (Isaacson & Scanziani, 2011). Although V1 neurons directly receive geniculate excitation, there is also intracortical excitation within V1 (Douglas et al., 1995). Moreover, what we estimate does not necessarily reflect the synaptic excitatory or inhibitory inputs a neuron directly receives. For example, a neuron could decrease its firing rate in response to light because its excitatory inputs are inhibited by light. Consequently, the estimated excitation and inhibition should best be interpreted as a measure of how a neuron’s response varies as a function of light and dark stimuli, and not simply as synaptic weighting.

Distinguishing excitation to dark from inhibition to light (and vice-versa) has been enabled by the use of rich stimuli such as natural images, combined with our simple model architecture. Had we attempted to make the model more complex and biologically realistic, the results we obtain from the analysis might be more dependent on the particular sort of model we use and thus become problematic to interpret. Natural image stimuli lead to more robust system identification than with synthetic stimuli (Talebi & Baker, 2012), and perhaps more importantly, they ensure that neurons simultaneously receive visual stimuli that both increase and decrease their firing rate in different parts of their receptive fields - this allows the machine learning algorithm to distinguish between excitation from one pathway and inhibition from the other.

### Dark-dominance due to weaker inhibition from dark stimuli

Dark-dominance in V1 has previously been thought to originate from relatively greater lateral geniculate excitation from the OFF pathway (Jin et al., 2008). However, recent findings suggest dark-dominance might instead be caused by stronger intracortical inhibition from light than dark stimuli (Taylor et al., 2018). At each neuron’s optimal latency, our results support the latter hypothesis by showing ON inhibition to be much stronger than OFF inhibition, while we do not find a significant difference between ON and OFF excitation.

These findings might help explain why dark-dominance is strongest in layer 2/3 of primate V1 (Yeh et al., 2009). If dark-dominance were principally due to stronger lateral geniculate excitation from the OFF pathway, we would expect dark-dominance to be at least as strong in layer 4 than in the other layers, since this is where most LGN neurons synapse. While two-thirds of the neurons in primate layer 4 show dark-dominance, this effect is much stronger in layers 2/3 where almost every neuron is dark-dominant (Yeh et al., 2009). This laminar difference might be due to pyramidal neurons in primate layers 2/3 receiving extensive inhibition, as has been shown in the mouse (Kätzel et al., 2011), with inhibition being stronger to light than dark stimuli (Taylor et al., 2018).

Since this study utilized recordings from polytrodes that did not extend across all the cortical layers, a laminar analysis was not feasible. A useful future direction could be to replicate this experiment with linear-array probes to obtain simultaneous recording across all V1 layers, to investigate the laminar dependence of dark-dominance.

### Time dynamics of dark-dominance

A novel finding of this study is how the dark-dominance changes as a function of latency. We observe the dark-dominance effect at the 0-13.3 and 13.3-26.7 latencies, but instead find a slight light-dominance at the 13.3-26.7 latency. We were able to find this relationship between latency and dark-dominance because we estimate and analyze light and dark responses at every latency for each neuron. Other studies have focused on each neuron’s optimal latency (e.g. Yeh et al., 2009), which still clearly shows the dark-dominance effect (Figure 3) but neglects the effect of latency on the strength of dark responses. Consequently, dark responses were thought to be on average stronger in general, whereas we find this effect to be specific to the earlier latencies.

This relationship between dark-dominance and latency should not be too surprising, considering dark-dominant V1 neurons have previously been found to respond 3-6 ms faster than light-dominant neurons (Komban et al., 2014). These faster dark responses in V1 have been attributed to faster OFF than ON LGN responses (Jin et al., 2008; Jin et al., 2011). While we do find the 0-13.3 ms latency to be dark-dominant due to stronger OFF than ON excitation (Figure 4B), most neurons have poor responses at this latency. The dark-dominance effect is most salient at the 13.3-26.7 ms latency, when response strength peaks and dark-dominance is due to weaker inhibition to dark stimuli (Figure 4C). These results are consistent with findings from Taylor et al. (2018), who found intracortical inhibition to be stronger for light than for dark stimuli. Therefore, we interpret the dark-dominance results at each neuron’s optimal time lag from Yeh et al. (2009) and Jansel et al. (2019) as mostly due to weaker inhibition rather than stronger excitation to dark stimuli.

### Relationship to orientation selectivity

This study also brings a new perspective on the intracortical mechanisms of orientation selectivity, and helps explain why V1 neurons are less orientation-selective in their early time lags (Ringach et al., 1997; Shapley et al., 2003). Due to the absence of direct inhibition from the LGN to V1 (Ferster & Lindström 1983, Martin & Whitteridge 1984; Montero, 1986), the lagged onset of orientation selectivity was previously attributed to the delay imposed by the necessity of intracortical inhibitory interneurons (Ringach et al., 1997; Shapley et al., 2003). We do find inhibition strength to be positively correlated with orientation selectivity (Figure 7D; see Li, Yang, Liang, Xia & Zhou, 2008). However, we also find neurons with higher orientation selectivity to have more balanced light/dark responses (Figure 7B). Consistent with these results, responses from 0 to 26.7 ms, which are lower in orientation selectivity (Figure 7A), are also biased towards dark stimuli (Figure 4A) and have stronger excitation than inhibition (Figure 7C). In contrast to the first two latencies, the 26.7-40 ms latency has high orientation selectivity (Figure 7A) and relatively balanced responses between light and dark stimuli (Figure 4A, 4D). Because both dark-dominance and stronger inhibition at the 13.3-26.7 latency are due to slower inhibition to dark stimuli (Figure 4C), the reason why neurons are less orientation selective at the 13.3-26.7 than at the 26.7-40 ms latency could possibly be due to this slower inhibition to dark stimuli.

### Conclusion

In conclusion, we use a novel machine learning approach to bring new insights to the phenomenon of stronger dark responses in visual cortex neurons. We find the dark-dominance effect to only occur in the early latencies, and to be due to slower inhibition to dark stimuli. We also show how weaker average inhibition to dark stimuli is related to weaker orientation selectivity in the early latencies. The questions of how and why primary visual cortex neurons receive slower inhibition to dark than to light stimuli, and whether these findings vary across laminae, could be fruitful subjects of future investigation.

## Acknowledgements

We wish to thank Guangxing Li for contributions to software and technical support in experiments, and Philippe Nguyen and Amol Gharat for their important contributions to data collection and spike-sorting. Funded by Canadian Institutes of Health Research (CIHR) grant MOP-119498 to C.B.

